# Asynchronous phylogeographic and demographic dynamics of rodent community in the low latitude Asia

**DOI:** 10.1101/2023.03.14.532549

**Authors:** Gaoming Liu, Cheng-Min Shi, Huajing Teng, Jian-Xu Zhang, Quansheng Liu

## Abstract

It is increasing evident that demographic history and phylogeographic consequences of past climate changes were unfolded locally and varied from region to region. Despite the high Murinae rodent species richness and endemism in the low latitude Asia, how the past climatic fluctuations shaped the phylogeographic and demographic history in this area remains unknown. Here we trapped 253 field Murine individuals and successfully amplified COI gene sequence for DNA barcoding. The phylogenetic tree showed the Murinae diversification included ten species belong to Rattini and Murini tribes. The divergence dating suggested that the most common ancestor (TMRCA) of each rodent species was estimated in Early or Middle Pleistocene. Bayesian skyline plot (BSP) exhibited the onset of population growth of seven Murinae rodents occurred at penultimate or last glaciation, and while *R. losea* and *R. norvegicus* keep effective population size constant through their elapsed time. Additionally, the six rodent species range of refugia area in the LGM projected by ecological niche models (ENMs) exhibited expander than the suitable area on present, meanwhile the remaining four rodent species showed contracted refugia regions. Hence, our results suggested that the rodent community displayed the asynchronous demographic and phylogeographic dynamics in the low latitude Asia.

## INTRODUCTION

It is increasing evident that climate has exerted pervasive influences on all hierarchy of biological systems, ranging from individuals, populations, communities to entire ecosystems. Biological impacts of climate changes have been particularly well-exemplified by a wealth of studies probing through the phylogeographic lens on the genetic variations of animals and plants during the Quaternary ice ages (Avise 2000, Parvizi *et al.* 2019, Rey-Iglesia *et al.* 2021). It appeared that the phylogeographic consequences of past climate changes were unfolded locally and varied from region to region, although Quaternary climate fluctuations were a global phenomenon. For example, Lessa et al. (2003) demonstrated that populations of four small mammals in the high latitude North America synchronously expanded during the late Quaternary, while populations of 11 small mammals in the low altitude Amazon were stable during the same period (Lessa *et al.* 2003). Carnaval *et al.* (2009) also showed similar findings on a more restricted geographic scale within the tropics. However, there were also evidences suggested that regional communities might have respond to past climate change idiosyncratically (Burbrink *et al.* 2016), and different taxonomic groups responded differently to past climatic changes (Antonelli *et al.* 2018).

The low latitude regions, especially tropics, are the geographical zone farthest from polar ice sheets, and have been little affected by glaciers formation processes during the glacial cycles of the Quaternary (Zong 2007). In comparison with temperate regions, low latitude regions have maintained greater climatic stability during the glacial ages. Additionally, low latitude regions harbor particularly high biodiversity and species endemic to the tropical regions (Tonkin *et al.* 2016). The low latitude Asia is one of centers of diversity and endemism of Murinae rodents, a sub-family of the Muridae family (Latinne et al. 2013). Murine is subdivided ten tribes based on the molecular identification, two of which were Rattini and Murini tribes (Lecompte *et al.* 2008). The former encompasses 35 genera and the latter harbors 2 genera (Musser & Carleton 2005). However, Murinae field identification remains difficult, presumably owing to multiple ambiguous effectors, including the limited interspecfic external morphological divergence and/or large scale interspecific morphological variations (e. g., *Rattus* amd *Niviventer* genus). Also, comparative phylogeographic approaches are relatively unexplored in the tropics and how the past climatic fluctuations in shaped phylogeographic history of tropical species remains uncertain.

In this study, we used DNA barcoding and phylogenetic analysis to delimit the Murine species in the low latitude Asia. We estimated the divergence time and Bayesian skyline plot (BSP) to show the demographic history of those rodents. We also constructed ecological niche models to project the suitable distributional areas of rodents in the present day and LGM.

## MATERIAL AND METHODS

### Sample collection and DNA sequencing

A total of 253 field individuals of Murine rats were trapped in Guangdong Province, China. All individuals were diagnosed to putative taxa based on morphology. Total genomic DNA was extracted from tissues of tail tips using a TailGen DNA extraction Kit (CWBIO, Beijing, China) or following a modified phenol-chloroform extraction procedure (Zhang, Hewitt 1998). We amplified and sequenced partial fragment of the mitochondrial cytochrome oxidase I subunit gene (mtCOI) for all of the 253 samples using primers BatL5310 (5’-CCTACTCRGCCATTTTACCTATG-3’) and R6036R (5’-ACTTCTGGGTGTCCAAAGAATCA-3’) (Robins *et al.* 2007). Polymerase chain reactions (PCRs) were carried out in volumes of 30 uL consisting of 1X reaction buffer, 1.5 mM of MgCl2, 0.2 mM of each dNTPs, 0.3 uM of each primer, 0.3 unit of Taq DNA polymerase (Promega, Shanghai, China) and ~40 ng of genomic DNA, with following thermal profile: an initial denaturation step at 94°C for 4min was followed by 35 cycles of at 94°C for 30 s, at 51 °C for 30 s and at 72°C for 40 s, and a final extension step at 72°C for 2 min. The purified PCR products were sequenced using the ABI PRISM BigDye Terminator V3.1 Cycle Sequencing Kit and sequences were resolved on an ABI 3730XL automated sequencer (Applied Biosystems, Foster City, CA, USA). All sequences are deposited in GenBank under accession numbers of xxxxxxxx-xxxxxxxx.

### Phylogenetic analysis

The new sequences were edited to remove unreliably resolved bases at both ends and aligned using the CLUSTAL X 2.0 program (Larkin *et al.* 2007). After checked the possible interference of mitochondrial pseudogenes following the recommendation of Zhang and Hewitt (Zhang & Hewitt 1996) and (Bensasson *et al.* 2001), the alignment was collapsed into haplotypes using DnaSP 5.10 (Librado, Rozas 2009). Using the unique haplotypes as queries, the standard nucleotide BLAST searches were performed in the NCBI database (https://blast.ncbi.nlm.nih.gov) to retrieve the homologous sequences. We downloaded the hit homologous sequences with query coverage of 100% and identity scores > 97%, a threshold slightly lower than the within species similarity found in Murinae species. As all BLAST hit species were belong to Murinae, three sequences for *Gerbillus* species (Gerbillnae) were included as outgroups based on study of Rowe et al. (2016). These sequences were aligned with the haplotypes generated in this study and subjected to phylogenetic analyses and divergence time estimation (Table 1; Table S1, Supporting Information).

**Table 1.** Sample statistics information and neutral test as well as the estimated time of ten rodent species

| Clade | SN | SNh | TN | TNh | Hd | $\pi$ (%) | Tajima's D | P-value | Fu's Fs | P-value | R<sup>2</sup> | P-value | Mean | 95%HPD |
| --- | --- | --- | --- | --- | --- | --- | --- | --- | --- | --- | --- | --- | --- | --- |
| *R. tanezumi* | 10 | 2 | 44 | 30 | 0.914 | 0.581 | -1.930 | 0.006** | -27.650 | 0.000*** | 0.040 | 0.000*** | 0.59 | 0.37-0.87 |
| *R. tiomanicus* | 1 | 1 | 22 | 17 | 0.965 | 0.878 | -1.445 | 0.051 | -8.069 | 0.003** | 0.075 | 0.020* | 1.11 | 0.68-1.62 |
| *R. losea* | 72 | 6 | 82 | 16 | 0.726 | 0.530 | -0.857 | 0.198 | -2.649 | 0.216 | 0.070 | 0.217 | 0.57 | 0.36-0.86 |
| *R. andamanensis* | 73 | 16 | 94 | 32 | 0.868 | 0.857 | -0.875 | 0.202 | -11.749 | 0.013* | 0.071 | 0.234 | 0.96 | 0.62-1.37 |
| *R. norvegicus* | 18 | 5 | 26 | 9 | 0.831 | 0.620 | 0.470 | 0.727 | -0.142 | 0.495 | 0.145 | 0.739 | 0.54 | 0.28-0.87 |
| *Ba. indica* | 8 | 2 | 17 | 10 | 0.875 | 0.814 | -1.009 | 0.152 | -1.459 | 0.237 | 0.102 | 0.120 | 0.79 | 0.46-1.19 |
| *N. fulvescens* | 38 | 3 | 60 | 24 | 0.728 | 1.203 | -1.481 | 0.044* | -3.852 | 0.150 | 0.059 | 0.037* | 0.93 | 0.58-1.35 |
| *N. confucianus* | 16 | 3 | 29 | 14 | 0.813 | 0.445 | -1.785 | 0.022* | -6.141 | 0.025* | 0.055 | 0.001*** | 0.65 | 0.38-1.01 |
| *Be. bowersi* | 1 | 1 | 9 | 7 | 0.917 | 1.396 | -1.052 | 0.151 | -0.003 | 0.414 | 0.195 | 0.803 | 0.60 | 0.30-0.98 |
| *M. caroli* | 16 | 8 | 18 | 10 | 0.915 | 0.795 | -1.559 | 0.045* | -1.296 | 0.259 | 0.100 | 0.131 | 1.25 | 0.75-1.82 |

Trapped sample number of individuals (SN), Trapped sample number of haplotypes (SNh), Toal number of individuals (N), Total number of haplotypes (Nh), haplo- type diversity (Hd), nucleotide diversity ( $\pi$ ), 95% highest posterior density (95% HPD), \* indicates $p < 0.05$ , \*\* indicates $p < 0.01$ , \*\*\* indicates $p < 0.001$ .

Phylogenetic analyses were performed using both maximum likelihood (ML) and Bayesian approaches with the best-fit model of nucleotide substitution identified by jModelTest 2 (Darriba *et al.* 2012). ML analysis was carried out with the software PhyML 3.0 (Guindon *et al.* 2010). Nodal support was assessed through 1000 bootstrap replicates. Bayesian analysis was carried out with MrBayes 3.2.6 (Ronquist *et al.* 2012). Analysis was initiated with random starting trees, and run for 5 × 10^6^ generations with four Markov chains. Trees were sampled every 500 generations and the first 25% of trees were discarded as burn-in.

### Population genetic analyses

Summary statistics of population genetics, the number of haplotypes (H), haplotype diversity (Hd), and nucleotide diversity (π), were calculated for each of the monophyletic lineages recognized in the phylogenetic analysis using DnaSP 5.10 (Librado & Rozas 2009). The pairwise distances between lineages/species were calculated based the Kimura’s two-parameter model (Kimura 1980) as implemented in MEGA version X (Kumar *et al.* 2018).

### Divergence time estimated

Divergence time and evolutionary rates for each branch were estimated with a relaxed molecular clock approach implemented in BEAST 1.8.4 (Drummond *et al.* 2012), as the strict molecular clock model was rejected by likelihood-ratio tests (2Δℓ = 492.66, d.f. = 162, P < 0.001). The time to the most recent common ancestor (*T_MRCA_*) and evolutionary rate for each lineage were estimated through a fossil-calibrated approach. Two fossil calibrations were used: (1) the split of *Mus/Rattus* (10.5-14.0 Ma) (dos Reis *et al.* 2012, Kimura *et al.* 2013, Kimura *et al.* 2015), for which we set a normal prior distribution with mean 12.2 Ma and standard deviation (SD) 0.9 Ma for the calibrating node; (2) the most recent common ancestor of the *Mus* (7.3-8.3Ma) (Kimura *et al.* 2015, Rowe *et al.* 2016), for which we used mean 7.8 Ma and SD 0.24 Ma for the normal prior distribution. The evolutionary rate change was explicitly modelled using uncorrelated lognormal distribution across trees. A Yule process prior was used for modelling speciation. One hundred million Markov chain Monte Carlo (MCMC) searches were performed and sampled every 10000 generations. Convergence of the MCMC chains was checked with Tracer 1.7 (Rambaut *et al.* 2018). Maximum clade credibility (MCC) tree, posterior means and 95% highest posterior densities (HPDs) of ages, and evolutionary rates were identified and annotated using TreeAnnotator 1.8.4 (Rambaut & Drummond 2015).

### Demographic history inferences

Historical changes in the effective population size were assessed using three approaches which use different information contents of DNA sequences. First, the Tajima’s *D* (Tajima 1989), *Fs* (Fu 1997) and *R2* (Ramos-Onsins, Rozas 2002) statistics were calculated. Tajima’s *D* test is based on the number of segregating sites; Fu’s *Fs* test use information from the haplotype distribution, while *R2* test use information of the mutation frequency. These statistics have high power to detect population expansion for nonrecombining regions of the genome under a variety of different circumstances, either when population sample sizes are large (*Fs*) or when sample sizes are small and the number of segregating sites is low (*R2*) (Ramos-Onsins, Rozas 2002). The significance of each test was assessed by generating null distributions from 10,000 coalescent simulations in DnaSP version 5.10 (Librado, Rozas 2009). Significantly large negative *D* (*P* < 0.05) and *Fs* (*P* < 0.02) values and significantly positive *R2* (*P* < 0.05) values were taken as evidence of a population expansion. Second, mismatch distribution using information from the distribution of the pairwise sequence differences was employed. The observed distribution of pairwise differences was compared to data simulated under the sudden and spatial expansion models. Multimodal distributions suggest a historically stable population size, whereas unimodal distributions imply population expansion (Rogers & Harpending 1992). Sum of square deviations (SSD) was used to assess the fit of the data to the model. The Harpending’s raggedness index quantifies the smoothness of the observed pairwise difference distribution with a non-significant (*P* > 0.05) result indicates an expanding population (Harpending 1994). Mismatch distribution were analyzed using Arlequin ver 3.5 (Excoffier & Lischer 2010). Third, changes in effective population size (*N_e_*) over time were explored by coalescent-based Bayesian skyline plot (BSP) (Drummond *et al.* 2005) using BEAST 1.8.4 (Drummond *et al.* 2012). Unlike above tests which are based on summary statistics, BSP make use of all the historical information contained within a sample of DNA sequences from a given population. The estimate of evolutionary rate for each of the lineage/species was used to calibrate BSP analysis using a strict molecular clock model. We ran the analysis for 5 × 10^8^ iterations, treating the first 10% iterations as burn-in, then subsequently sampling genealogies and model parameters every 5000 generations post burn-in.

### Ecological niche modeling

We constructed ecological niche models (ENMs) to check for spatial changes of suitable distributional areas of rats. Geographic coordinates for the samples used in the demographic analysis were combined to the occurrence localities of the respective species obtained from the Global Biodiversity Information Facility (GBIF, www.gbif.org). We masked the focus region to an area (85~150 °E, - 10~40 °N) reasonably larger than the Southeast Asia (85~150 °E, −10~40 °N) to examine historical range shifts. The location points downloaded from the GIBF were cleaned up by excluding the duplications and possible mislabeled coordinates, and were further filtered by keeping only one occurrence point in each of 1 × 1 degree cells to reduce the effects of sampling bias (Table S2, Supporting Information). To estimate habitat suitability for each species, we used the bioclimatic variables at 2.5-arcminute resolution from the WorldClim (www.worldclim.org/) database. Climatic layers were cropped to the above masked region to avoid sampling unrealistic background data inflating the strength of predictions (Elith *et al.* 2011). We excluded highly correlated climatic variables with variance inflation factor (VIF) > 10 (Naimi *et al.* 2014), to avoid over-fitting of the model and hampering interpretability of results. The maximum entropy algorithm implemented in MaxEnt version 3.4.1 was used to model the species distribution (Phillips *et al.* 2006, Phillips *et al.* 2017). The present-day ENMs were projected to the past conditions during LGM (21,000 years ago) simulated using the Community Climate System Model (CCSM) available at the WorldClim website. MaxEnt was run with a convergence threshold of 10^-5^ (default) and maximum number of iterations of 10, 000 for 10 replicates with cross-validate. Model performance was assessed via the area under the ROC (receiver operating characteristic) curve (AUC) statistic. We employed the 10 percent training presence threshold (10%TP) to create binary map of predictions. The 10%TP is conservative, which under-predicts known range by excluding the 10% most extreme observations. Predictions based on this threshold reflect the core of the present species range (Morueta-Holme *et al.* 2010).

## RESULTS

### Phylogeny and diversity of rodent community

A total of 253 field Murine individuals were trapped and COI gene was successfully amplified and sequenced, collapsed into 47 unique haplotypes. Blasted best hit results for the NCBI database showed those haplotypes represented ten putative species in five genera. Sample size for each classified species inferred from the combined blast best hit results and phylogenetic relationship varied from 1 to 73 field Murinae individuals (Table 1). Combined best hit COI sequences from the GenBank database, we finally obtained 393 sequence alignments and collapsed into 168 haplotypes used for the phylogenetic inference.

The phylogenetic relationship of all unique haplotypes reconstructed by PhyML and MrBays were congruent with each other and the topologies were highly supported with ten rodent species (Figure1). The results were consisted with blasted best hit putative species. Obviously, phylogenetic ingroup first divergent two tribes: Rattini and Murini, the former harbored *Rattus tenezumi, R. tiomanicus, R. losea, R. andamanensis, R. norvegicus, Bandicota indica, Niviventer fulvescens, N. confucianus and Berylyms bowersi* meanwhile the latter tribe included *Mus caroli* and *M. parahi* (Figure 1). Additionally, *Niviventer* genus is endemic to Southeast Asia (Musser & Carleton, 2005).

**Figure 1.**
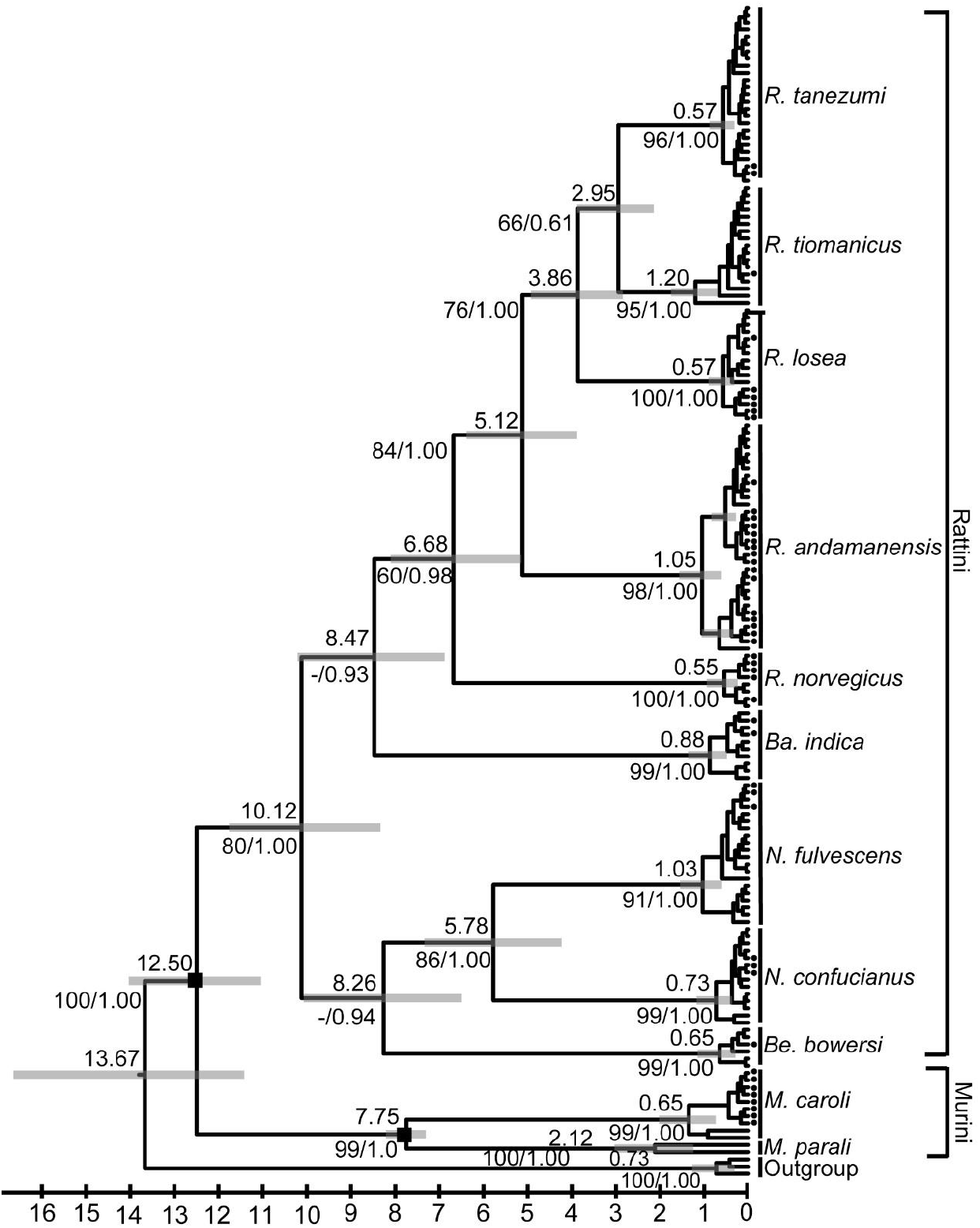
Phylogenetic relationship and divergence dating of Muridae based on mitochondrial COI gene. The estimated mean divergence time are showed above branches. Gray bar represents 95% highest posterior density. Numbers below the branches indicate Bayesian posterior probabilities and maximum likelihood bootstrap support values. The black quadrangle indicates the fossil calibration point. The tip with black dot indicates the trapped sample haplotypes.

The nucleate and haplotype diversity of each delimitated species varied range from 0.530% to 1.396% and from 0.726 to 0.965, respectively (Table 1). To evaluate the divergent levels between pairwise species, we calculated the genetic distances, the maximum genetic distance occurred between *N. fulvescens* and *Ba. Indica,* was 0.194. Conversely, the minimum was 0.067 between congeneric *R. timanicus* and *R. tanezumi,* which were reciprocal monophyletic group (Figure 1; Table 2).

**Table 2.** The pairwise genetic distance of the rodent community

|  | *M. caroli* | *Ba. indica* | *Be. bowersi* | *N. confucianus* | *N. fulvescens* | *R. norvegicus* | *R. andamanensis* | *R. losea* | *R. tiomanicus* | *R. tanezum* |
| --- | --- | --- | --- | --- | --- | --- | --- | --- | --- | --- |
| *M. caroli* |  |  |  |  |  |  |  |  |  |  |
| *Ba. indica* | 0.178 |  |  |  |  |  |  |  |  |  |
| *Be. bowersi* | 0.169 | 0.174 |  |  |  |  |  |  |  |  |
| *N. confucianus* | 0.813 | 0.181 | 0.148 |  |  |  |  |  |  |  |
| *N. fulvescens* | 0.170 | 0.200 | 0.156 | 0.126 |  |  |  |  |  |  |
| *R. norvegicus* | 0.189 | 0.159 | 0.145 | 0.161 | 0.181 |  |  |  |  |  |
| *R. andamanensis* | 0.185 | 0.161 | 0.153 | 0.181 | 0.168 | 0.132 |  |  |  |  |
| *R. losea* | 0.180 | 0.166 | 0.163 | 0.173 | 0.177 | 0.114 | 0.108 |  |  |  |
| *R. tiomanicus* | 0.172 | 0.178 | 0.144 | 0.164 | 0.165 | 0.120 | 0.099 | 0.087 |  |  |
| *R. tanezum* | 0.177 | 0.167 | 0.152 | 0.164 | 0.159 | 0.119 | 0.104 | 0.079 | 0.067 |  |

### Divergence time of rodent community

We estimated the divergence time under the uncorrected lognormal relaxed clock. Figure 1 showed that the first divergence in Rattini occurred in Middle/Late Miocene, corresponding to 9.87Ma (95% highest posterior density, 95% HPD: 8.13-11.57Ma), generating the two clades, both divergent in Late Miocene, specifically at 8.33Ma (95% HPD: 6.65-10.08Ma) and 8.08Ma (95% HPD: 6.42-9.94Ma). The estimated time of the most recent common ancestor (TMRCA) of each rodent species was occurred in Early or Middle Pleistocene, thereinto TMRCA of *M. caroli* was the most ancient at 1.25Ma (95% HPD: 0.75-1.82Ma) and meanwhile *R. losea* was most recent at 0.57Ma (95% HPD: 0.36-0.86Ma) (Table 1).

### Asynchronous demographic histories of rodent community

Significant signs of population expansion were detected in six of the ten studied species by at least one of the three tests (Table 1). Values of Tajima’s *D* and Fu’s *Fs* for all species are negative, except for *N. norvigicus.* Values for *R2* are all small, four of which are significant (*R. tanezumi, R. tanezumi, N. fulvescens* and *N. confucianus;* Table 1). Only *R. tanezumi* and *N. confucianus* in all three tests were showed significant (Table 1).

In neutrality tests, Tajima’s *D* value were significantly negative, which showed population may experience expansion. In an extensive variety of cases, the behavior of Fu’s *Fs* and R2 test held more powerful than Tajima’s *D* for detecting population growth (Ramos-Onsins, Rozas 2002). Tajima’s *D* and Fu’s *Fs* tests with significantly negative and R2 test with significant for *R. tanezumi* and *N. confucianus* suggested recent population expansion (Table 1). For *N. fulvescnes* population, Tajima’s *D* and R2 values were both significant, indicating population underwent population expansion. Fu’s *Fs* test with significantly negative and R2 test with significantly positive for *R. tiomanicus* showed a wave of population expansion (Table 1). For *M. caroli* population, only significantly negative Tajima’s *D* value, indicating a conceivable scenario of population growth (Table 1). Fu’s *Fs* value was negative and significant for *R. andamensis* suggested the population increase scenario (Table 1).

BSP showed *R. tanezumi* experienced population growth through approximately 0.15-0.03Ma, and *R. tuimanicus* and *N. fulvescens* population had a sudden increase the effective size from about 0.20Ma until now (Figure 2). Although *R. andamensis* likewise underwent a sudden population expansion, they occurred at rough 0.075Ma until now (Figure 2). For *Ba. indica* and *N. confulvescnes* population, both had a gradual population increase through their demographic history, yet began at different point-in-time, aroud 0.28Ma and 0.18Ma, respectively (Figure 2). *Be. bowersi* and *M. caroli* population went through similar historical demography, initial population stable size then a small amplitude increases at approximately 0.15Ma (Figure 2). Nevertheless, *R. losea* and *R. norvegicus* population held steady size through their elasped time (Figure 2). Hence, BSPs displayed that seven species experienced population expansion and meanwhile two species kept the invariable effective size, which suggested the asynchronous demographic histories.

**Figure 2.**
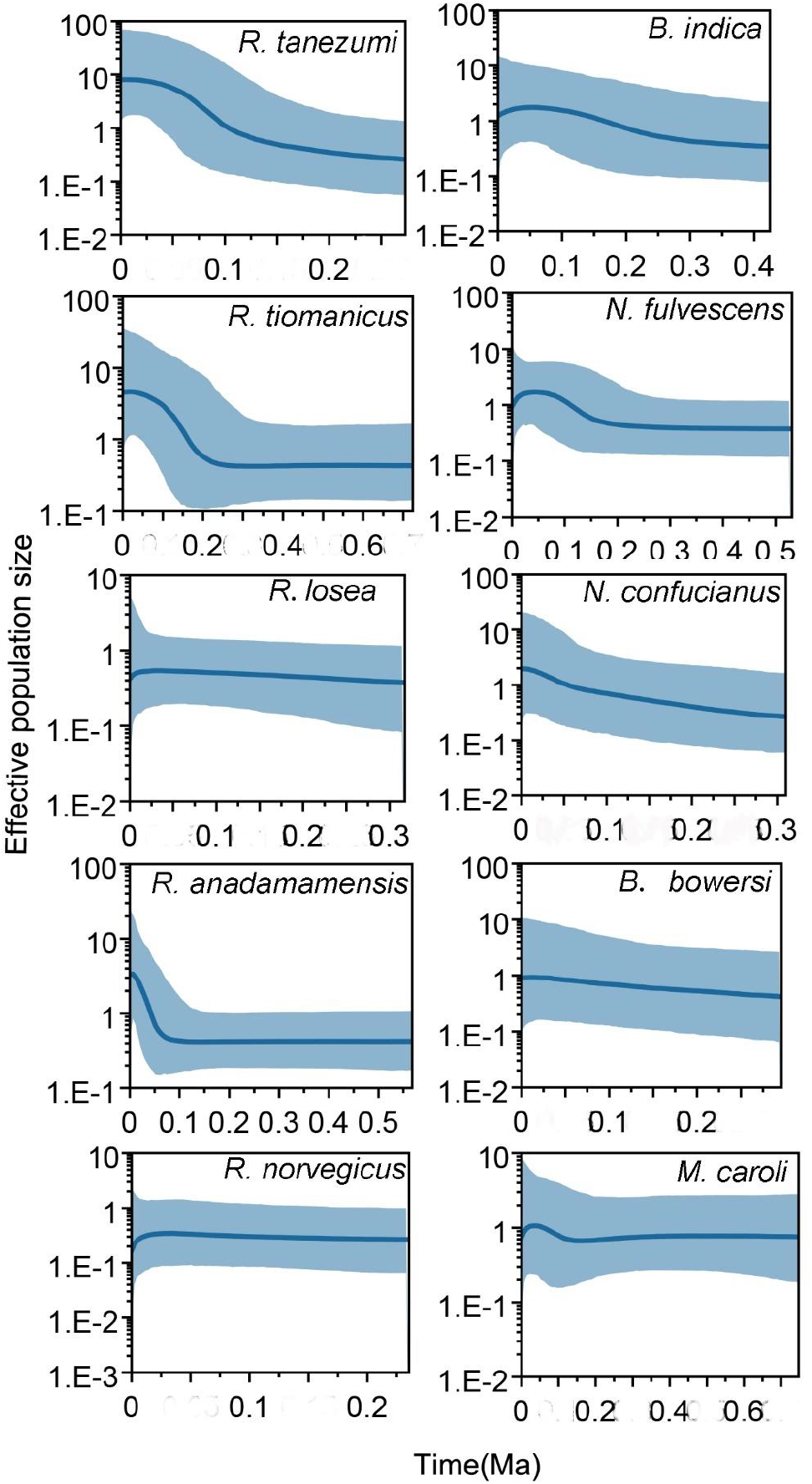
Bayesian skyline plot of historical demography for rat community in the low latitude Asia. The solid line represents the median numbers of the effective population sizes. The gray shade areas correspond to 95% highest posterior densities (95% HPDs).

### Asynchronous phylogeographic dynamics of rat community over the Last Glacial Maximum

We obtained 8 bioclimatic variables excluding highly correlated climatic variables, including mean diurnal range (BIO2), isothermality (BIO3), mean temperature of wettest quarter (BIO8), precipitation of wettest month (BIO13), precipitation of driest month (BIO14), precipitation seasonality (BIO15), precipitation of warmest quarter (BIO18) and precipitation of coldest quareter (BIO19).

Visualizations of ENMs are shown in Figure 3. Projections onto the climatic conditions of present day revealed the suitable area was major in low latitude regions (Figure 3). There are six species showed larger refugia area and more south regional boundaries over the LGM, including *Be. bowersi, N. confucianus, N. fulvescens, R. andamensis, R. tanezumi and R. tiomanicus,* compared with the present-day suitable area. And meanwhile the remaining four rodent refugia areas were shrunk (Figure 3). Hence, the rodent species showed the asynchronous phylogeographic dynamics over the LGM. The overlap regions of the projected suitable areas under two climate scenarios for each rodent species was almost through the Tropic of Cancer (Table 3; Figure 3), then extended southward and northward, exclude *R. timonanicus* only located southern regions of the Tropic of Cancer (Figure 3).

**Figure 3.**
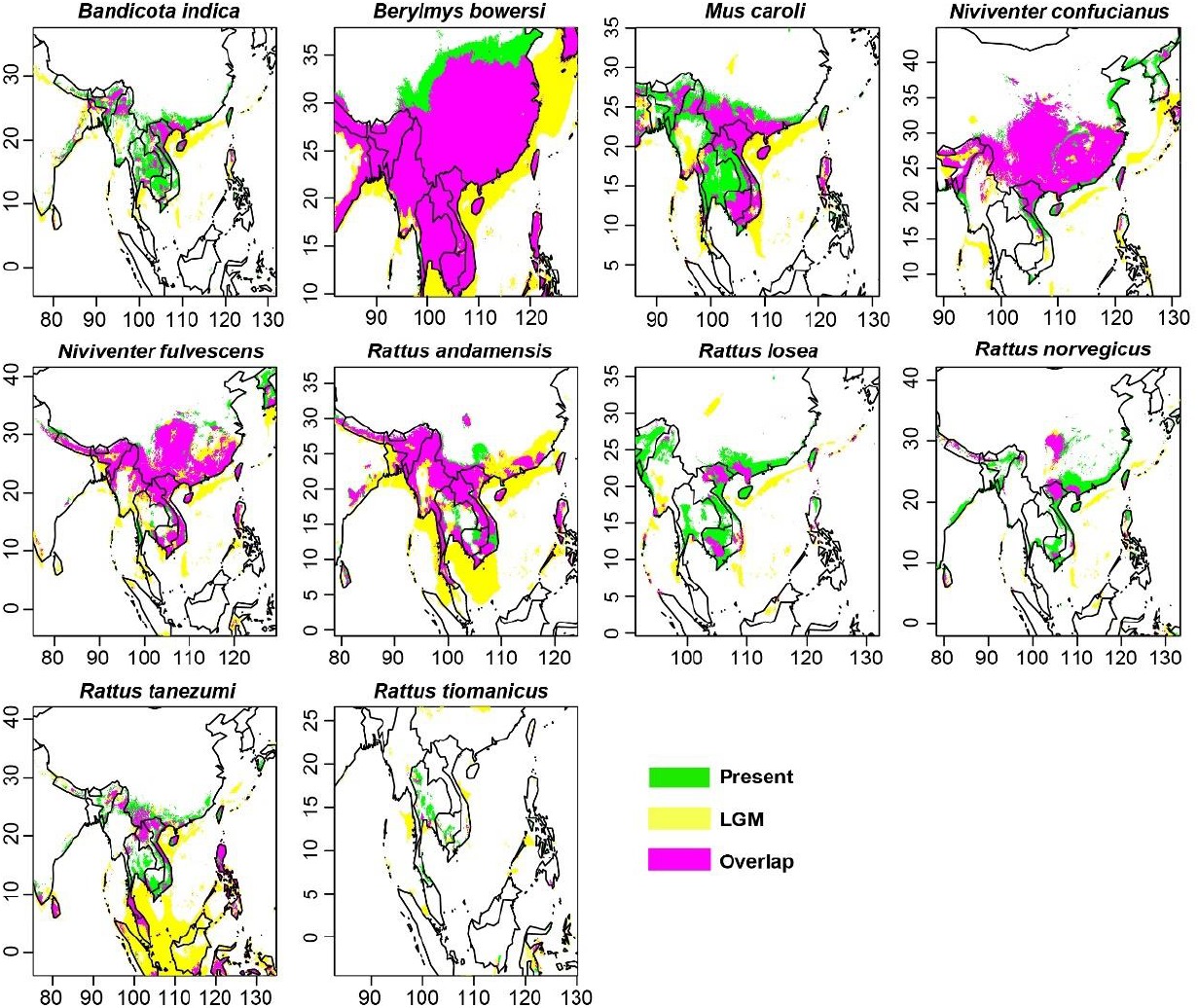
The projection of the suitable area (green) in the present day and refugia areas (yellow) in the LGM as well as overlap of both areas (pink) for the rate community using the ENMs.

**Table 3.** The projection area in the present and LGM bioclimate variables for the NEMs

| Species | Present | LGM | Overlap | Overlap/Present (%) | Overlap/LGM (%) |
| --- | --- | --- | --- | --- | --- |
| *Ba. indica* | 95560 | 84818 | 36855 | 38.57 | 43.45 |
| *Be. bowersi* | 331686 | 387299 | 302206 | 91.11 | 78.03 |
| *M. caroli* | 135211 | 126576 | 75979 | 56.19 | 60.03 |
| *N. cofucianus* | 176736 | 182996 | 144110 | 81.54 | 78.75 |
| *N. fulvescens* | 176272 | 228188 | 149018 | 84.54 | 65.30 |
| *R. andamensis* | 127894 | 228629 | 111789 | 87.41 | 48.90 |
| *R. losea* | 54958 | 28943 | 11086 | 20.17 | 38.30 |
| *R. norvegicus* | 83632 | 42043 | 24280 | 29.03 | 57.75 |
| *R. tanezumi* | 113203 | 232044 | 64968 | 57.39 | 27.99 |
| *R. tiomanicus* | 34231 | 44599 | 12471 | 36.43 | 27.96 |

## DISCUSSION

The phylogenetic tree demonstrated the trapped 253 field Murine individuals classified ten species in five genera, belonging to two tribes: Rattini and Murini (Figure 1). Murinae rats were difficult to demarcate due to conspicuous biodiversity and special evolutionary characteristics, such as rapid adaptive radiation and short divergence time (Conroy, Cook 1999, Steppan *et al.* 2004, Steppan *et al.* 2005, Robins *et al.* 2007, Rowe *et al.* 2011).

Sample size for each classified species inferred from the combined blast best hit results and phylogenetic relationship varied from 1 to 73 field Murinae individuals (Table 1). Singletons, such as *R. tiomanicus* and *Be. Bowersi,* were not discarded, as in most cases, a single typical specimen could still provide a reliable benchmark as an *ad hoc* method for species delimitation in DNA barcoding (Lim *et al.* 2012, Li *et al.* 2015). We don’t deny that singleton was merely obtained from one sample rodent, the dependability needed more field individuals to intensify.

In the phylogenetic tree, the reciprocal monophyletic group between *R. tanezumi* and *R. tiomanicus* was not well-supported. Although both not difficult confused the external morphology, *R. tiomanicus* always had whitish underparts and sleeker upperparts with less promient guard hairs, compared with the *R. tanezumi,* (Francis, Barrett 2008). In this study, *R. tiomanicus* samples from Robins *et al.* (2007) (Table S1, Supporting Information) were recognized as *R. tanezumi* by taxomonic characteristics, and samples from Latinne *et al.* (2013) were identified clusters *Rattus* R3 lineage, which was phylogenetic species delimitated by Pages *et al.* (2010) according to General Mixed Yule Coalescnet (GMYC) model delimited species boundaries. Balakirev & Rozhnov (2012) combined morphological data and biogeography and proposed a new system of nomination claimed that phylogenetic R3 was *R. tiomanicus.* Apart from *Rattus* genus, *Niviventer* genus was endemic to Southeast Asia (Musser & Carleton, 2005) and one of taxonomically ambiguous members of the Murinae (Musser, Carleton 2005, Balakirev *et al.* 2011, He, Jiang 2015). The samples of *N. ling* (corresponding to *N. sp.* in Table S1, Supporting Information) is morphologically distinct from *N. fulvescnes* in head, body and tail length, but it still clustered *N. fulvesnces* group as an operational taxonomic unit (OTU) in phylogenetic tree (Lu *et al.* 2015). Similarly, *N. confucianus* group was highly phylogenetic diversity (Zhang *et al.* 2016) and included 27 OTUs, one of which was *N. lotipes* (corresponding to *N. sp.* in Table S1, Supporting Information) (Lu *et al.* 2015). The phylogenetic analyses of previous studies have found the *Be. bowersi* can divided into two highly divergent lineages, “Ber2” lineage (Latinne *et al.* 2013) or “Be2” lineage (Pages *et al.* 2010). In this study, we no evidenced these two lineages. We cannot deny that there was merely one sample of *Be. bowersi,* which maybe not enough to powerful support the species delimitation for this sample. We need more evidence, such as samples field capture and/or taxonomic characteristics.

Divergence time showed that TMRCA of all Murinae species located in low latitude Asia was estimated in Middle/Late Pleistocene (Figure 1). To test the accuracy of estimated time for the species, we compare the estimated date with the known fossil record. The ancestor of *N. confucianus,* considered as *N. preconfucianus,* was found at Zhoukoudian, Beijing in the early Pleistocene (approximate 1.20 Ma), which is older than our estimated split time of *N. confucianus* (95% HPD: 0.38-1.01Ma) (Table 1). The most recent fossil record of *N. confucianus* was from Panxian Dadong, Guizhou Province, China and occurred in middle-late Pleistocene (0.13-0.30 Ma) (Schepartz *et al.* 2003, Bekken *et al.* 2004). Hence, the split of *N. confucianus* samples in south China was occurred at 0.35-0.96 Ma, which was covered by fossil records (0.13-1.20 Ma). Another species *R. norvegicus,* originated in northern China and Mongolia, has enjoyed a worldwide distribution (Lin *et al.* 2012, Puckett *et al.* 2016). The most representative ancient fossils of this species were in Southwestern in China, about 1.20-1.60Ma (Jin *et al.* 2008, Yuan *et al.* 2009), then found recent fossils in Guangdong Province in late Middle Pleistocene (Wu, Wang 2012). Hence the fossil time beyond the estimated time of TMRCA for *R. norvegicus* range from 0.29Ma to 0.88Ma (Table 1). Additionally, our estimated TMRCA of *M. carilo* was more ancient than previous study estimated (Shimada *et al.* 2007), which maybe result from divergence time estimated methods were different.

BSP displayed that the episode of population expansion for six Murinae specie (*R. tanezumi, R. tiomanicus*, *Ba. indica*, *N. fulvescens*, *N. confulvescens*, *Be. bowersi* and *M. caroli*) began from approximate 0.28-0.15Ma (Figure 2), accordance to the penultimate glaciation equivalent to marine isotope stage (MIS) 6-8 (Yi *et al.* 2005, Zhao *et al.* 2011). The onset of the population growth for *R. andamanensis* was nearly consist with the first stage of last glaciation (Yi *et al.* 2005, Zhao *et al.* 2011). However, *R. losea* and *R. norvegicus* population held steady size through their elapsed time. Overall, Pleistocene climate, especially glaciation, modeled the population dynamics of Murinae species in low latitude Asia, but the population demographic history of rodent community respond to glaciation was asynchronous.

The six rodent species range of refugia area in the LGM projected by NEMs exhibited expander than the suitable area on present, meanwhile the remaining four rodent species showed contracted refugia regions (Table 3; Figure 3). Hence, the phylogeographic dynamics of rodent community in the low latitude Asia was asynchronous over the LGM. This refugia area expansion scenario was inconsistent with the median and high latitude hypothesis, which refugia area in the LGM showed contacted regions compared with suitable area in the present (Avise 2000, Thomé *et al.* 2010, Zhao *et al.* 2019).

Eustatic sea levels fell by up to 120m below the present levels due to the sequestering of ice during major glacial periods, resulted in the repeated formation of land bridges between formerly isolated land masses (Lambeck *et al.* 2002), such as between Asian mainland and Hainain, Sumatra, Java and Borneo (Voris 2000, Woodruff 2010). The present continental shelf was exposed to become land in the glaciation, which providing the new suitable area for the rodent community. All island samples in *Niviventer* and *Rattus* species can immigrate through the land bridges during the episode of population expansion in the penultimate and last glaciation. All rodent community recolonized the exposed land to refugia even though refugia area was shrunk for some rodent species (Figure 3). The phenomenon of the taxa expanded the exposed new land over the LGM was also found in the low latitude Neotropical forest (Leite *et al.* 2016). Therefore, the refugia area dynamics of the low latitude Asian rodent community over the LGM is asynchronous notwithstanding on the synchronous benefit of the new suitable exposed land and greater climatic stability.

Mitochondrial data in the study displayed no distinguishingly increase effective size for *R. losea* and *R. norvegicus* to support the dispersal scenario (Figure 2), but the matrilineal inheritance cannot reveal the history of the species as a whole. Indeed, sex-biased dispersals have been reported in some mice (Van Hooft *et al.* 2008, Solmsen *et al.* 2011). For example, the nuclear single nucleotide polymorphisms (SNPs) of *R. norvegicus* illuminated the range across all over the world (Puckett *et al.* 2016) not the confined range showed in figure 3. Therefore, the mitochondrial sequences only supply the maternal inheritance history for effective population size and suitable areas respond to the glaciation climate and environment, not show the paternal genetic history.

## Supporting information

Table S1 Information regarding the sequences included in this study.

Table S2 The location of rodent species

## AUTHOR’S CONTRIBUTIONS

Quansheng Liu trapped the rat samples and extracted the DNA from the samples. Gaoming Liu amplified the PCR and the bioinformatic analyses as well as submitted the sequences to GenBank, along with wrote the manuscript with input from the other authors. Cheng-Min Shi provided some guides for the evolutionary analyses. Huajing Teng and Jian-Xu Zhang designed the framework of this study. Quansheng Liu and Cheng-Min Shi revised the manuscript. All authors read and approved the final version of the manuscript.

## ACKNOWLEDGEMENTS

This project was supported by the National Natural Science Foundation of China (grant No. 32170455), the State Key Laboratory of Integrated Management of Pest Insects and Rodents (IPM2005) and by the State Key Laboratory of North China Crop Improvement and Regulation (YJ2020028).

## COMPENTING INTERESTS

The authors declare that they have no competing interests.

## Supporting Information

Table S1 Information regarding the sequences included in this study.

Table S2 The location information used for the ENMs

