## Supplementary material for "Asynchronous phylogeographic and demographic dynamics of rodent community in the low latitude Asia": Table S1 Information regarding the sequences included in this study.

| Clade | Species | Accension No. | Location | Coordinates | Reference |
|---|---|---|---|---|---|
| *M. caroli* | *M. caroli* | KJ530557<br>KJ530558 | | | (Tsangaras *et al.* 2014) |
| *M. pahari* | *M. pahari* | JF445030<br>JF445031 | Quang Nam, Ngoc Linh Base Camp, Vietnam | | |
| *Be. bowersi* | *Be. bowersi* | JN105107<br>JN105105<br>JN105106<br>JN105104<br>JN105108 | Son La province, Vietnam | 104.77° E, 21.14° N; 104.77° E, 21.14° N | (Balakirev *et al.* 2011) |
| | | HM217542<br>HM217580 | Loei, Thailand | | (Pages *et al.* 2010) |
| *N. confucianus* | *N. confucianus* | HM031911<br>HM031912<br>HM031913 | Wuzhishan, Hainan, China | | (Lu *et al.* 2012) |
| | *N. sp.* | KF739944<br>KF739945 | Wuyishan, Fujiang, China | 118.02° E, 27.46° N | (Lu *et al.* 2015) |
| | | KF739946 | Yingtan, Jiangxi, China | 117° E, 28.33° N | |
| | | KF739948<br>KF739951 | Qingyuan, Zhejiang, China | 118.80° E, 27.80° N | |
| | | KF739949 | Wuxi, Jiangsu, China | 120.30° E, 31.5° N | |
| | | KF739950 | Wuxi, Jiangsu, China | 120.30° E, 31.57° N | |
| | | KF739947 | Yingtan, Jiangxi, China | 117.0° E, 28.23° N | |
| | | KF739953 | Xingyi, Guizhou, China | 104.89° E, 25.09° N | |
| | | KF739952 | Libo, Guizhou, China | 107.88° E, 25.42° N | |
| *N. fulvescens* | *N. fulvescens* | HM031915<br>HM031918<br>HM031924<br>HM031925<br>HM031927<br>HM031928 | Hainan, China | | (Lu *et al.* 2012) |
| | | KR868707 | Emei Moutain, Sichuan Province, China | 103.33° E, 29.55° N | (Yong *et al.* 2016) |
| | | KP455488 | | | |
| | *N. sp.* | KF739970<br>KF739971 | Lincang Snow Mountain, Lincang, Yunnan, China | 100.23° E, 23.90° N | (Lu *et al.* 2015) |
| | | KF739966<br>KF739967 | Wuyishan, Fujiang, China | 118.02° E, 27.46° N | |
| | | KF739969 | Qingyuan, Zhejiang, China | 118.80° E 27.80° N | |

| | | | | | |
|---|---|---|---|---|---|
| | | KF739968 | Tengchong, Yunnan, China | 98.47° E,<br>25.03° N | |
| | | KF739964 | Xingshan, Hubei, China | 110.50° E,<br>31.56° N | |
| | | KF739965<br>KF739963 | Gongga Mountain, Sichuan, China | 101.50° E,<br>29.50° N | |
| | | KF739961 | Chishui, Guizhou, China | 105.42° E,<br>28.34° N | |
| *Ba. indica* | *Ba. indica* | FR775844 | Vinh Long, Vietnam | | (Balakirev & Rozhnov 2012) |
| | | HM217509 | Ratchaburi, Thailand | | (Pages *et al.* 2010) |
| | | HM217517 | Nakhon Ratchasima, Thailand | | |
| | | HM217554 | Phrae, Thailand | | |
| | | HM217574 | Loei, Thailand | | |
| | | JQ307468 | India | | |
| | | JQ362480 | India | | |
| | | JQ362481 | India | | |
| | | KT029807 | Guiping, Guangxi Province, China | 110.03° E,<br>23.05° N | (Wang *et al.* 2016) |
| *R. norvegicus* | *R. norvegicus* | EF186576<br>EF186577 | Raiatea, Society Islands, French Polynesia | | (Robins *et al.* 2007) |
| | | KM577634 | | | |
| | | M27315 | | | (Pepe *et al.* 1983) |
| | | X14848 | | | (Grosskopf & Feldmann 1981) |
| | | KP100657 | | | |
| | | KP233827 | | | |
| *R. andamanensis* | *R. andamanensis* | JQ793911<br>JQ793916<br>JQ793917 | Ruili, Yunnan, China | | (Lu *et al.* 2012) |
| | | JQ793910<br>JQ793912<br>JQ733913<br>JQ793914 | Lincang, Yunnan, China | | |
| | | KC010293<br>KC010294 | Nan, Thailand | 100.60° E,<br>19.38° N | (Latinne *et al.* 2013) |
| | | HM031791<br>HM031798<br>HM031802<br>HM031804<br>HM031805<br>HM031810<br>HM031811<br>HM031815<br>HM031818<br>HM031820<br>HM031827<br>HM031835 | Qiongzhou, Hainan, China<br>Wuzhishan, Hainan, China<br>Chengmai, Hainan, China | | (Lu *et al.* 2012) |

| | | | | | |
|---|---|---|---|---|---|
| *R. losea* | *R. losea* | HM031880<br>HM031881<br>HM031882<br>HM031884<br>HM031888<br>HM031889<br>HM031891<br>HM031892<br>HM031895<br>HM031896 | Qiongzhou, Hainan, China<br>Chengmai, Hainan, China | | (Lu *et al.* 2012) |
| *R. tiomanicus* | *R. sp.* | EF186621 | Yogyakarta, Indonesia | | (Robins *et al.* 2007) |
| | | EF186614<br>EF186616 | Jakarta, Indonesia | | |
| | | EF186626<br>EF186627 | Northern Sulawesi, Indonesia | | |
| | | KC010277 | Chaiyaphum, Thailand | 101.5667° E, 16.5667° N | (Latinne *et al.* 2013) |
| | | KC010278<br>KC010284 | Nakhon Ratchasima, Thailand | 101.2333° E, 14.5833° N | |
| | | KC010279<br>KC010280<br>KC010283 | Lopburi, Thailand | 100.8667° E, 14.8° N | |
| | | KC010281 | Petchaburi, Thailand | 99.9333° E, 14.8° N | |
| | | KC010282 | Petchaburi, Thailand | 99.7333° E, 13.7167° N | |
| | | KC010285<br>KC010286 | Petchaburi, Thailand | 99.7667° E, 13.6333° N | |
| | | KC010287 | Kanchanaburi, Thailand | 99.5667° E, 13.9667° N | |
| | *R. tiomanicus* | FR775809 | Nam Cat Tien, Dong Nai province, Vietnam | | (Balakirev & Rozhnov 2012) |
| | | FR775816<br>FR775817<br>FR775819 | Vinh Long, Vietnam | | |
| *R. tanezumi* | *R. tanezumi* | EF186620<br>EF186622 | Yogyakarta, Indonesia | | (Robins *et al.* 2007) |
| | | EF186625 | Amami Island, Japan | | |
| | | EF186606 | Jakarta, Indonesia | | |
| | | EU273712 | | | |
| | | KC010257 | Nan, Thailand | 100.6833° E 18.45° N | (Latinne *et al.* 2013) |
| | | KC010258 | Nan, Thailand | 100.75° E 19.1667° N | |
| | | KC010259 | Kanchanaburi, Thailand | 99.5667° E 13.9667° N | |
| | | KC010260 | Prachuap Khiri Khan, Thailand | 99.8833° E 12.1° N | |

| | | | | |
|---|---|---|---|---|
| | | KC010261<br>KC010262 | Nan, Thailand | 100.60° E,<br>19.38° N |
| | | KC010263 | Saraburi, Thailand | 101.13° E,<br>14.63° N |
| | | KC010264 | Petchabun, Thailand | 100.93° E,<br>15.77° N |
| | | KC010265<br>KC010266 | Krabi, Thailand | 98.87° E,<br>8.12° N |
| | | KC010267 | Krabi, Thailand | 98.82° E,<br>8.60° N |
| | | KC010268 | Nakhon Si Thammarat, Thailand | 99.67° E,<br>8.18° N |
| | | KC010269 | Petchabun, Thailand | 101.03° E,<br>15.73° N |
| | | KC010270 | Prachuap Khiri Khan, Thailand | 99.90° E,<br>11.42° N |
| | | KC010271 | Petchaburi, Thailand | 99.93° E,<br>12.85° N |
| | | KC010272 | Petchaburi, Thailand | 99.92° E,<br>13.10° N |
| | | KC010273 | Petchaburi, Thailand | 99.95° E,<br>12.85° N |
| | | KC010274 | Chumphon, Thailand | 99.07° E,<br>10.45° N |
| | | KC010275 | Nakhon Ratchasima, Thailand | 101.22° E,<br>14.6° N |
| | | KC010276 | Nakhon Sawan, Thailand | 100.43° E<br>15.28° N |
| | | KF011916 | | |
| | | JQ906931 | | |
| Outgroup | *G. nanus* | KF422708<br>KF422713<br>KF722714 | | |


Balakirev AE, Abramov AV, Rozhnov VV (2011) Taxonomic revision of Niviventer (Rodentia, Muridae) from Vietnam: a morphological and molecular approach. *Russian Journal of Theriology* **10**, 1-26.

Balakirev AE, Rozhnov VV (2012) Contribution to the species composition and taxonomic status of some Rattus inhabiting southern Vietnam and Sundaland. *Russian Journal of Theriology* **11**, 33-45.

Grosskopf R, Feldmann H (1981) Analysis of a DNA segment from rat-liver mitochondria containing the genes for the cytochrome-oxidase subunit-I, subunit-II and subunit-III, atpase subunit-6, and several transfer-RNA genes. *Current Genetics* **4**, 151-158.

Latinne A, Waengsothorn S, Rojanadilok P*, et al.* (2013) Diversity and endemism of Murinae rodents in Thai limestone karsts. *Systematics and Biodiversity* **11**, 323-344.

Lu L, Chesters D, Zhang W*, et al.* (2012) Small mammal investigation in spotted fever focus with DNA-barcoding and taxonomic implications on rodents species from Hainan of China. *PLoS One* **7**, e43479.

Lu L, Ge D, Chesters D*, et al.* (2015) Molecular phylogeny and the underestimated species diversity of the endemic white-bellied rat (Rodentia: Muridae:Niviventer) in Southeast Asia and China. *Zoologica Scripta* **44**, 475-494.

Pages M, Chaval Y, Herbreteau V*, et al.* (2010) Revisiting the taxonomy of the Rattini tribe: a phylogeny-based delimitation of species boundaries. *BMC Evolutionary Biology* **10**, 27.

Pepe G, Holtrop M, Gadaleta G*, et al.* (1983) Non random patterns of nucleotide substitutions and codon strategy in the mammalian mitochondrial genes-coding for identified and unidentified reading frames. *Biochemistry International* **6**, 553-563.

Robins JH, Hingston M, Matisoo-Smith E, Ross HA (2007) Identifying Rattus species using mitochondrial DNA. *Molecular Ecology Notes* **7**, 717-729.

Tsangaras K, Wales N, Sicheritz-Ponten T*, et al.* (2014) Hybridization Capture Using Short PCR Products Enriches Small Genomes by Capturing Flanking Sequences (CapFlank). *PLoS One* **9**, 10.

Wang SH, Cong HY, Kong LM, Motokawa M, Li YC (2016) Complete mitochondrial genome of the greater bandicoot rat Bandicota indica (Rodentia: Muridae). *Mitochondrial DNA Part A* **27**, 4349-4350.

Yong B, Wei H, Jia Q, Chen S (2016) Sequencing and analysis of complete mitochondrial genome of Niviventer fulvescens (Muridae). *Mitochondrial DNA* **27**, 3650-3651.
