## Supplementary material for "Asynchronous phylogeographic and demographic dynamics of rodent community in the low latitude Asia": Table S2 The location of rodent species

| species | lon | lat |
|---|---|---|
| Bandicota_indica | 121.076 | 22.77124 |
| Bandicota_indica | 120.2522 | 23.05729 |
| Bandicota_indica | 121.0566 | 24.92047 |
| Bandicota_indica | 120.6107 | 24.40384 |
| Bandicota_indica | 104.8118 | 11.47137 |
| Bandicota_indica | 105.131 | 12.024 |
| Bandicota_indica | 105.28 | 17.67 |
| Bandicota_indica | 106.057 | 20.637 |
| Bandicota_indica | 91.3965 | 23.0107 |
| Bandicota_indica | 91.1822 | 23.4565 |
| Bandicota_indica | 102.97 | 13.56 |
| Bandicota_indica | 102.2905 | 20.11723 |
| Bandicota_indica | 101.7 | 19.3 |
| Bandicota_indica | 97.62 | 16.49 |
| Bandicota_indica | 96.8838 | 20.6036 |
| Bandicota_indica | 99.17 | 18.67 |
| Bandicota_indica | 95.2213 | 18.8215 |
| Bandicota_indica | 104.8314 | 22.7694 |
| Bandicota_indica | 106.463 | 20.696 |
| Bandicota_indica | 108.3968 | 11.2108 |
| Bandicota_indica | 108.4874 | 11.2005 |
| Bandicota_indica | 106.71 | 15.34 |
| Bandicota_indica | 106.1278 | 14.6306 |
| Bandicota_indica | 102.02 | 14.6083 |
| Bandicota_indica | 102.0167 | 14.36667 |
| Bandicota_indica | 113.8808 | 22.24806 |
| Bandicota_indica | 113.8833 | 22.2333 |
| Bandicota_indica | 110.8333 | -7.58333 |
| Bandicota_indica | 108.3333 | -6.46667 |
| Bandicota_indica | 100.47 | 5.5 |
| Bandicota_indica | 100.37 | 6.08 |
| Bandicota_indica | 106.5333 | -6.1 |
| Bandicota_indica | 106.8833 | -6.1 |
| Bandicota_indica | 108.25 | 16.02 |
| Bandicota_indica | 85.31667 | 27.71667 |
| Bandicota_indica | 85.3 | 27.73333 |
| Bandicota_indica | 91.9 | 25.66667 |
| Bandicota_indica | 91.76667 | 25.46667 |
| Bandicota_indica | 97 | 25.36667 |
| Bandicota_indica | 102.1 | 21.68333 |
| Bandicota_indica | 96.08 | 21.97 |
| Bandicota_indica | 96.015 | 17.125 |
| Bandicota_indica | 85.33333 | 19.76667 |
| Bandicota_indica | 110.0333 | 23.05 |
| Berylmys_bowersi | 101.372 | -1.70866 |
| Berylmys_bowersi | 101.89 | 20.51 |
| Berylmys_bowersi | 102.18 | 19.76 |
| Berylmys_bowersi | 105.6419 | 21.45389 |
| Berylmys_bowersi | 101.8349 | 3.1032 |
| Berylmys_bowersi | 101.7386 | 3.71634 |
| Berylmys_bowersi | 91.76667 | 25.46667 |
| Berylmys_bowersi | 94.05 | 25.3 |
| Berylmys_bowersi | 97.71667 | 27.9 |
| Berylmys_bowersi | 97.7 | 28.11667 |
| Berylmys_bowersi | 103.8667 | 22.35 |
| Berylmys_bowersi | 110 | 24 |
| Berylmys_bowersi | 119.3667 | 25.71667 |

| | | |
|---|---|---|
| Berylmys_bowersi | 104.0419 | 20.4169 |
| Berylmys_bowersi | 104.7688 | 21.1395 |
| Berylmys_bowersi | 104.772 | 21.1363 |
| Berylmys_bowersi | 99.3667 | 8.5 |
| Mus_caroli | 121.2487 | 25.01075 |
| Mus_caroli | 120.349 | 23.27571 |
| Mus_caroli | 102.7833 | 17.66203 |
| Mus_caroli | 105.28 | 17.67 |
| Mus_caroli | 102.1 | 21.68333 |
| Mus_caroli | 102.2905 | 20.1172 |
| Mus_caroli | 103.73 | 13.43 |
| Mus_caroli | 102.97 | 13.56 |
| Mus_caroli | 96.015 | 17.125 |
| Mus_caroli | 97.646 | 16.2825 |
| Mus_caroli | 103.0667 | 21.66666 |
| Mus_caroli | 104.1134 | 20.459 |
| Mus_caroli | 121.0585 | 16.9177 |
| Mus_caroli | 106.5105 | 10.9966 |
| Mus_caroli | 102.61 | 17.97 |
| Mus_caroli | 101.8833 | 17.1 |
| Mus_caroli | 103.4833 | 22.11666 |
| Mus_caroli | 103.0246 | 24.92754 |
| Mus_caroli | 98.53817 | 3.21196 |
| Mus_caroli | 115.3833 | -8.55 |
| Mus_caroli | 112.9455 | -8.01419 |
| Mus_caroli | 99.06667 | 19.38333 |
| Mus_caroli | 98.91667 | 18.91667 |
| Mus_caroli | 100.5709 | 13.84833 |
| Mus_caroli | 100.9167 | 14.51667 |
| Mus_caroli | 113.0333 | 22.53333 |
| Mus_caroli | 113.1167 | 23.03333 |
| Mus_caroli | 108.5 | 12.08333 |
| Mus_caroli | 108.35 | 11.68333 |
| Mus_caroli | 109.5476 | 19.40766 |
| Mus_caroli | 109.5667 | 19.51667 |
| Mus_caroli | 106.71 | 15.34 |
| Mus_caroli | 128.1667 | -3.68333 |
| Mus_caroli | 129.8333 | -4.51667 |
| Mus_caroli | 127.2 | -8.08333 |
| Mus_caroli | 120.4667 | -8.58333 |
| Mus_caroli | 122.65 | -8.55 |
| Niviventer_confucianus | 103.6667 | 32.5 |
| Niviventer_confucianus | 103.3671 | 34.60283 |
| Niviventer_confucianus | 105.897 | 16.9507 |
| Niviventer_confucianus | 105.13 | 17.75 |
| Niviventer_confucianus | 105.97 | 23.12 |
| Niviventer_confucianus | 107.88 | 25.48 |
| Niviventer_confucianus | 108.3 | 29.41667 |
| Niviventer_confucianus | 107.5 | 29.65 |
| Niviventer_confucianus | 104.2333 | 28.75 |
| Niviventer_confucianus | 103.3003 | 29.10803 |
| Niviventer_confucianus | 102.6833 | 25.1 |
| Niviventer_confucianus | 103.8667 | 22.35 |
| Niviventer_confucianus | 113.1 | 29.38333 |
| Niviventer_confucianus | 111.03 | 26.42 |
| Niviventer_confucianus | 119.3667 | 25.71667 |
| Niviventer_confucianus | 121.2667 | 24.15 |
| Niviventer_confucianus | 122.1167 | 48.55 |

| | | |
|---|---|---|
| Niviventer_confucianus | 98.48333 | 18.58333 |
| Niviventer_confucianus | 118.1667 | 26.63333 |
| Niviventer_confucianus | 119.3333 | 26.08333 |
| Niviventer_confucianus | 156.625 | -7.7917 |
| Niviventer_confucianus | 157.165 | -8.3083 |
| Niviventer_confucianus | 107.7668 | 33.95004 |
| Niviventer_confucianus | 107.3833 | 33.8805 |
| Niviventer_confucianus | 113.5667 | 37.33333 |
| Niviventer_confucianus | 101.45 | 43.5219 |
| Niviventer_confucianus | 118.02 | 27.46 |
| Niviventer_confucianus | 117 | 28.3333 |
| Niviventer_confucianus | 118.8 | 27.8 |
| Niviventer_confucianus | 117 | 28.2333 |
| Niviventer_confucianus | 104.89 | 25.09 |
| Niviventer_confucianus | 107.88 | 25.42 |
| Niviventer_fulvescens | 113.8833 | 22.2333 |
| Niviventer_fulvescens | 113.9042 | 22.23 |
| Niviventer_fulvescens | 101.3833 | 4.51667 |
| Niviventer_fulvescens | 100.2664 | 5.425813 |
| Niviventer_fulvescens | 105.13 | 17.75 |
| Niviventer_fulvescens | 105.21 | 17.72 |
| Niviventer_fulvescens | 107.02 | -6.75 |
| Niviventer_fulvescens | 106.707 | -6.7305 |
| Niviventer_fulvescens | 101.372 | -1.70866 |
| Niviventer_fulvescens | 101.3701 | -1.70853 |
| Niviventer_fulvescens | 108.35 | 11.68333 |
| Niviventer_fulvescens | 107.42 | 11.42 |
| Niviventer_fulvescens | 106.134 | 17.0534 |
| Niviventer_fulvescens | 106.0488 | 16.9574 |
| Niviventer_fulvescens | 108.0833 | 29.5 |
| Niviventer_fulvescens | 107.16 | 28.23 |
| Niviventer_fulvescens | 108.5 | 12.08333 |
| Niviventer_fulvescens | 106.3333 | 15.33333 |
| Niviventer_fulvescens | 107.89 | 21.85 |
| Niviventer_fulvescens | 105.97 | 23.12 |
| Niviventer_fulvescens | 103.54 | 11.83 |
| Niviventer_fulvescens | 96.4724 | 21.9883 |
| Niviventer_fulvescens | 95.7 | 26 |
| Niviventer_fulvescens | 97.50681 | 25.58487 |
| Niviventer_fulvescens | 102.8167 | 22.38333 |
| Niviventer_fulvescens | 102.9167 | 22.21666 |
| Niviventer_fulvescens | 97.71667 | 27.9 |
| Niviventer_fulvescens | 96.87 | 27.25 |
| Niviventer_fulvescens | 102.1 | 21.68333 |
| Niviventer_fulvescens | 99.13 | 20.05 |
| Niviventer_fulvescens | 109.85 | 29.18 |
| Niviventer_fulvescens | 110.17 | 28.67 |
| Niviventer_fulvescens | 98.91667 | 14.91667 |
| Niviventer_fulvescens | 101.15 | 14.57 |
| Niviventer_fulvescens | 85.53333 | 28 |
| Niviventer_fulvescens | 87.36692 | 27.39172 |
| Niviventer_fulvescens | 110.45 | -7.48 |
| Niviventer_fulvescens | 110.45 | -7.5 |
| Niviventer_fulvescens | 87.9333 | 27.00108 |
| Niviventer_fulvescens | 88.21667 | 27.01667 |
| Niviventer_fulvescens | 91.7 | 25.3 |
| Niviventer_fulvescens | 91.9 | 25.66667 |
| Niviventer_fulvescens | 93 | 26 |

| | | |
|---|---|---|
| Niviventer_fulvescens | 94.05 | 25.3 |
| Niviventer_fulvescens | 118.1667 | 26.63333 |
| Niviventer_fulvescens | 119.3667 | 25.71667 |
| Niviventer_fulvescens | 103.1 | 30.03333 |
| Niviventer_fulvescens | 103.3333 | 29.08333 |
| Niviventer_fulvescens | 88.71667 | 27.23333 |
| Niviventer_fulvescens | 102.25 | 30.25 |
| Niviventer_fulvescens | 110 | 24 |
| Niviventer_fulvescens | 109.5667 | 19.51667 |
| Niviventer_fulvescens | 109.5476 | 19.40766 |
| Niviventer_fulvescens | 118.096 | 31.0897 |
| Niviventer_fulvescens | 99.27 | 18.5 |
| Niviventer_fulvescens | 103.328 | 29.545 |
| Niviventer_fulvescens | 101.7833 | 17.0833 |
| Niviventer_fulvescens | 101.8667 | 16.5667 |
| Niviventer_fulvescens | 100.23 | 23.9 |
| Niviventer_fulvescens | 118.02 | 27.46 |
| Niviventer_fulvescens | 118.8 | 27.8 |
| Niviventer_fulvescens | 98.467 | 25.033 |
| Niviventer_fulvescens | 110.5 | 31.56 |
| Niviventer_fulvescens | 101.5 | 29.5 |
| Niviventer_fulvescens | 105.42 | 28.34 |
| Rattus_andamanensis | 113.9042 | 22.23 |
| Rattus_andamanensis | 114.0833 | 22.5 |
| Rattus_andamanensis | 92.4553 | 21.0958 |
| Rattus_andamanensis | 105.28 | 17.67 |
| Rattus_andamanensis | 105.05 | 17.84 |
| Rattus_andamanensis | 107.23 | 12.51 |
| Rattus_andamanensis | 107.45 | 11.12 |
| Rattus_andamanensis | 103.541 | 11.827 |
| Rattus_andamanensis | 103.54 | 11.83 |
| Rattus_andamanensis | 96.5635 | 22.0675 |
| Rattus_andamanensis | 96.4724 | 21.9883 |
| Rattus_andamanensis | 96.2344 | 17.7478 |
| Rattus_andamanensis | 97.0994 | 17.4442 |
| Rattus_andamanensis | 104.05 | 10.61667 |
| Rattus_andamanensis | 108.03 | 15.2 |
| Rattus_andamanensis | 105.6 | 22.42 |
| Rattus_andamanensis | 105.42 | 22.33 |
| Rattus_andamanensis | 107.7 | 12.87 |
| Rattus_andamanensis | 97.81667 | 27.08333 |
| Rattus_andamanensis | 96.9 | 27.2 |
| Rattus_andamanensis | 101.3 | 21.88333 |
| Rattus_andamanensis | 101.3333 | 21.95 |
| Rattus_andamanensis | 102.18 | 13.1083 |
| Rattus_andamanensis | 99.27 | 18.5 |
| Rattus_andamanensis | 91.9 | 25.66667 |
| Rattus_andamanensis | 91.76667 | 25.46667 |
| Rattus_andamanensis | 91.7 | 25.3 |
| Rattus_andamanensis | 90.33333 | 25.45 |
| Rattus_andamanensis | 88.71667 | 27.23333 |
| Rattus_andamanensis | 94.05 | 25.3 |
| Rattus_andamanensis | 95.55059 | 25.18472 |
| Rattus_andamanensis | 106.3333 | 15.33333 |
| Rattus_andamanensis | 102.9167 | 22.21666 |
| Rattus_andamanensis | 103.35 | 22.53333 |
| Rattus_andamanensis | 101.8333 | 21.51667 |
| Rattus_andamanensis | 109.5667 | 19.51667 |

| | | |
|---|---|---|
| Rattus_andamanensis | 92.78333 | 9.166667 |
| Rattus_andamanensis | 104.0419 | 20.4169 |
| Rattus_andamanensis | 100.6 | 19.3833 |
| Rattus_losea | 120.4057 | 23.70777 |
| Rattus_losea | 120.742 | 23.76544 |
| Rattus_losea | 121.4775 | 22.67433 |
| Rattus_losea | 120.6119 | 22.40506 |
| Rattus_losea | 120.9382 | 24.70527 |
| Rattus_losea | 121.6835 | 24.71445 |
| Rattus_losea | 118.3014 | 24.46292 |
| Rattus_losea | 119.6167 | 23.43333 |
| Rattus_losea | 119.9935 | 26.23329 |
| Rattus_losea | 106.1973 | 16.954 |
| Rattus_losea | 104.8375 | 17.5611 |
| Rattus_losea | 105.12 | 17.76 |
| Rattus_losea | 107.88 | 25.48 |
| Rattus_losea | 104.82 | 12.14 |
| Rattus_losea | 105.46 | 11.98 |
| Rattus_losea | 101.618 | 17.427 |
| Rattus_losea | 101.6714 | 17.4847 |
| Rattus_losea | 103.09 | 13.28 |
| Rattus_losea | 103.13 | 13.05 |
| Rattus_losea | 103.73 | 13.43 |
| Rattus_losea | 103.75 | 13.38 |
| Rattus_losea | 104.78 | 10.97 |
| Rattus_losea | 102.45 | 18.93 |
| Rattus_losea | 102.61 | 17.97 |
| Rattus_losea | 106.5105 | 10.9966 |
| Rattus_losea | 106.6407 | 10.938 |
| Rattus_losea | 105.547 | 21.263 |
| Rattus_losea | 105.6419 | 21.45389 |
| Rattus_losea | 104.6806 | 18.0083 |
| Rattus_losea | 104.4778 | 18.0736 |
| Rattus_losea | 102.45 | 14.7 |
| Rattus_losea | 116.1845 | 24.13319 |
| Rattus_losea | 109.5667 | 19.51667 |
| Rattus_losea | 109.5476 | 19.40766 |
| Rattus_losea | 105.764 | 21.16 |
| Rattus_norvegicus | 105.3304 | 12.3299 |
| Rattus_norvegicus | 100.5352 | 13.75139 |
| Rattus_norvegicus | 120.4145 | 23.60352 |
| Rattus_norvegicus | 120.5147 | 23.47731 |
| Rattus_norvegicus | 114.0168 | 22.36059 |
| Rattus_norvegicus | 113.302 | 23.1861 |
| Rattus_norvegicus | 108.22 | 16.05 |
| Rattus_norvegicus | 109.5119 | 18.25285 |
| Rattus_norvegicus | 119.3787 | 47.00723 |
| Rattus_norvegicus | 119.1612 | 47.10265 |
| Rattus_norvegicus | 123.0607 | 0.537206 |
| Rattus_norvegicus | 124.8282 | 1.34908 |
| Rattus_norvegicus | 101.8641 | 2.845213 |
| Rattus_norvegicus | 105.11 | 10.48 |
| Rattus_norvegicus | 106.088 | 9.63 |
| Rattus_norvegicus | 121.4467 | 31.20194 |
| Rattus_norvegicus | 118.7833 | 32.06667 |
| Rattus_norvegicus | 139.0046 | 36.01694 |
| Rattus_norvegicus | 139.1021 | 36.10571 |
| Rattus_norvegicus | 155 | 40 |

| | | |
|---|---|---|
| Rattus_norvegicus | 172.9076 | 52.90128 |
| Rattus_norvegicus | 173.1581 | 52.80878 |
| Rattus_norvegicus | 121.39 | 53.32 |
| Rattus_norvegicus | 122.4 | 53.46 |
| Rattus_norvegicus | 150.86 | 59.73 |
| Rattus_norvegicus | 150.8667 | 59.73333 |
| Rattus_norvegicus | 101.89 | 20.51 |
| Rattus_norvegicus | 106.057 | 20.637 |
| Rattus_norvegicus | 108.3968 | 11.2108 |
| Rattus_norvegicus | 108.4874 | 11.2005 |
| Rattus_norvegicus | 149.783 | 45.8585 |
| Rattus_norvegicus | 149.7832 | 45.8585 |
| Rattus_norvegicus | 150.141 | 46.08367 |
| Rattus_norvegicus | 150.164 | 46.09183 |
| Rattus_norvegicus | 120.5644 | 15.1908 |
| Rattus_norvegicus | 120.6166 | 16.3832 |
| Rattus_norvegicus | 132.2331 | 34.14444 |
| Rattus_norvegicus | 127.0683 | 37.56527 |
| Rattus_norvegicus | 112.73 | -7.28 |
| Rattus_norvegicus | 106.8833 | -6.1 |
| Rattus_norvegicus | 141.0236 | 42.62083 |
| Rattus_norvegicus | 141.4297 | 41.40194 |
| Rattus_norvegicus | 128.1597 | 26.73306 |
| Rattus_norvegicus | 127.7933 | 26.31639 |
| Rattus_norvegicus | 123.2667 | 9.3 |
| Rattus_norvegicus | 127.2833 | 45.36667 |
| Rattus_norvegicus | 126.9333 | 47.45 |
| Rattus_norvegicus | 103.6167 | 28.61667 |
| Rattus_norvegicus | 100.2 | 27.05 |
| Rattus_norvegicus | 107.986 | 29.3812 |
| Rattus_norvegicus | 108.1 | 29.29837 |
| Rattus_norvegicus | 88.35 | 22.58333 |
| Rattus_norvegicus | 119.6667 | -8.65 |
| Rattus_norvegicus | 124.5333 | -8.23333 |
| Rattus_norvegicus | 171.027 | 7.14641 |
| Rattus_tanezumi | 120.3291 | 22.78615 |
| Rattus_tanezumi | 121.4615 | 23.79894 |
| Rattus_tanezumi | 121.554 | 25.027 |
| Rattus_tanezumi | 121.5433 | 23.89241 |
| Rattus_tanezumi | 118.7333 | 9.7333 |
| Rattus_tanezumi | 121 | 10.8167 |
| Rattus_tanezumi | 141.5719 | -9.5169 |
| Rattus_tanezumi | 122.1352 | 18.54444 |
| Rattus_tanezumi | 121.0427 | 14.6511 |
| Rattus_tanezumi | 126.2167 | 6.95 |
| Rattus_tanezumi | 125.2667 | 6.9834 |
| Rattus_tanezumi | 101.6081 | 3.10697 |
| Rattus_tanezumi | 103.9143 | 1.315601 |
| Rattus_tanezumi | 114.38 | 38.31 |
| Rattus_tanezumi | 145.758 | 18.125 |
| Rattus_tanezumi | 145.7481 | 15.1925 |
| Rattus_tanezumi | 124.8834 | 12.15 |
| Rattus_tanezumi | 123.1167 | 9.5834 |
| Rattus_tanezumi | 107.88 | 25.48 |
| Rattus_tanezumi | 124.2333 | -9.53333 |
| Rattus_tanezumi | 124.12 | -9.97 |
| Rattus_tanezumi | 105.97 | 23.12 |
| Rattus_tanezumi | 103.35 | 22.53333 |

| | | |
|---|---|---|
| Rattus_tanezumi | 101.6889 | 17.49 |
| Rattus_tanezumi | 104.0933 | 17.0397 |
| Rattus_tanezumi | 96.015 | 17.125 |
| Rattus_tanezumi | 99.84927 | 16.0037 |
| Rattus_tanezumi | 100.6168 | 13.76732 |
| Rattus_tanezumi | 104.05 | 10.61667 |
| Rattus_tanezumi | 102.7224 | 24.74225 |
| Rattus_tanezumi | 103.2574 | 24.68003 |
| Rattus_tanezumi | 110.7514 | 19.62083 |
| Rattus_tanezumi | 109.7405 | 19.21959 |
| Rattus_tanezumi | 134.28 | 7.28 |
| Rattus_tanezumi | 129.5 | -3.25 |
| Rattus_tanezumi | 127.3333 | -0.8 |
| Rattus_tanezumi | 133.05 | -5.6 |
| Rattus_tanezumi | 116.68 | 6 |
| Rattus_tanezumi | 117.1 | 7.283333 |
| Rattus_tanezumi | 160 | -9.75 |
| Rattus_tanezumi | 127.68 | 0.88 |
| Rattus_tanezumi | 127.93 | 0.97 |
| Rattus_tanezumi | 136.0333 | -0.55 |
| Rattus_tanezumi | 134.0815 | -0.86704 |
| Rattus_tanezumi | 122.0333 | -8.23333 |
| Rattus_tanezumi | 119.35 | -9.43333 |
| Rattus_tanezumi | 106.92 | -6.88 |
| Rattus_tanezumi | 106.8 | -6.4 |
| Rattus_tanezumi | 110.4526 | -7.48758 |
| Rattus_tanezumi | 112.43 | -8.17 |
| Rattus_tanezumi | 98.6 | 10.15 |
| Rattus_tanezumi | 92.75 | 11.66667 |
| Rattus_tanezumi | 92.61667 | 10.76667 |
| Rattus_tanezumi | 119.93 | -1.03 |
| Rattus_tanezumi | 119.95 | -1.23 |
| Rattus_tanezumi | 98.58 | 3.5 |
| Rattus_tanezumi | 99.55 | 3.2 |
| Rattus_tanezumi | 110.68 | -1.08 |
| Rattus_tanezumi | 105.7 | -5 |
| Rattus_tanezumi | 85.19597 | 27.87583 |
| Rattus_tanezumi | 85.31667 | 27.71667 |
| Rattus_tanezumi | 93.5022 | 8.1128 |
| Rattus_tanezumi | 99.56667 | 9.216667 |
| Rattus_tanezumi | 145.7593 | 13.44143 |
| Rattus_tanezumi | 138 | -5 |
| Rattus_tanezumi | 140.7252 | -2.52273 |
| Rattus_tanezumi | 118.7833 | 32.06667 |
| Rattus_tanezumi | 94.43333 | 23.73333 |
| Rattus_tanezumi | 100 | 25.66667 |
| Rattus_tanezumi | 94.1543 | 23.9902 |
| Rattus_tanezumi | 108.3968 | 11.21077 |
| Rattus_tanezumi | 100.6833 | 18.45 |
| Rattus_tanezumi | 100.75 | 19.1667 |
| Rattus_tanezumi | 99.5667 | 13.9667 |
| Rattus_tanezumi | 99.8833 | 12.1 |
| Rattus_tanezumi | 100.6 | 19.3833 |
| Rattus_tanezumi | 101.1333 | 14.6333 |
| Rattus_tanezumi | 100.9333 | 15.7167 |
| Rattus_tanezumi | 98.8667 | 8.1167 |
| Rattus_tanezumi | 98.8167 | 8.6 |
| Rattus_tanezumi | 99.6667 | 8.1833 |

| | | |
|---|---|---|
| Rattus_tanezumi | 101.0333 | 15.7333 |
| Rattus_tanezumi | 99.9 | 11.4167 |
| Rattus_tanezumi | 99.9333 | 12.85 |
| Rattus_tanezumi | 99.9167 | 13.1 |
| Rattus_tanezumi | 99.95 | 12.85 |
| Rattus_tanezumi | 99.0667 | 10.45 |
| Rattus_tanezumi | 101.2167 | 14.6 |
| Rattus_tanezumi | 100.4333 | 15.2833 |
| Rattus_tiomanicus | 108.45 | -7.2 |
| Rattus_tiomanicus | 106.75 | -6.56667 |
| Rattus_tiomanicus | 110.4526 | -7.48758 |
| Rattus_tiomanicus | 110.3534 | -7.49632 |
| Rattus_tiomanicus | 114.2418 | -8.10512 |
| Rattus_tiomanicus | 113.0515 | 2.653333 |
| Rattus_tiomanicus | 113.051 | 2.6533 |
| Rattus_tiomanicus | 119.18 | 10.05 |
| Rattus_tiomanicus | 117.67 | 8.8 |
| Rattus_tiomanicus | 117.3 | 7.066667 |
| Rattus_tiomanicus | 116.55 | 6.08333 |
| Rattus_tiomanicus | 101.3 | -0.58333 |
| Rattus_tiomanicus | 101.3858 | -1.70356 |
| Rattus_tiomanicus | 105.4 | -6.1 |
| Rattus_tiomanicus | 104.5333 | -6.43333 |
| Rattus_tiomanicus | 102.4167 | 3.45 |
| Rattus_tiomanicus | 101.675 | 3.2 |
| Rattus_tiomanicus | 104.1667 | 2.78333 |
| Rattus_tiomanicus | 103.8476 | 1.88144 |
| Rattus_tiomanicus | 120.2 | 12 |
| Rattus_tiomanicus | 119.9 | 12.3 |
| Rattus_tiomanicus | 104.2667 | -4.03333 |
| Rattus_tiomanicus | 104.5333 | -4.41667 |
| Rattus_tiomanicus | 106.7992 | -6.12723 |
| Rattus_tiomanicus | 106.55 | -6.5 |
| Rattus_tiomanicus | 114.8333 | -3.41667 |
| Rattus_tiomanicus | 115.4167 | -2.63333 |
| Rattus_tiomanicus | 101.3833 | 4.51667 |
| Rattus_tiomanicus | 100.2458 | 5.4875 |
| Rattus_tiomanicus | 100.1 | 3.98 |
| Rattus_tiomanicus | 110.3333 | 1.48333 |
| Rattus_tiomanicus | 118.55 | 5.53333 |
| Rattus_tiomanicus | 115.45 | 3.75 |
| Rattus_tiomanicus | 115.5167 | 3.75 |
| Rattus_tiomanicus | 97.88333 | 4.8 |
| Rattus_tiomanicus | 117.1 | 8.2334 |
| Rattus_tiomanicus | 105.75 | -6.96667 |
| Rattus_tiomanicus | 116.15 | -8.6333 |
| Rattus_tiomanicus | 92.81667 | 11.11667 |
| Rattus_tiomanicus | 101.5667 | 16.5667 |
| Rattus_tiomanicus | 101.2333 | 14.5833 |
| Rattus_tiomanicus | 100.8667 | 14.8 |
| Rattus_tiomanicus | 99.9333 | 14.8 |
| Rattus_tiomanicus | 99.7333 | 13.7167 |
| Rattus_tiomanicus | 99.7667 | 13.6333 |
| Rattus_tiomanicus | 99.5667 | 13.9667 |
